# Tertiary structural motif sequence statistics enable facile prediction and design of peptides that bind anti-apoptotic Bfl-1 and Mcl-1

**DOI:** 10.1101/425926

**Authors:** Vincent Frappier, Justin M. Jenson, Jianfu Zhou, Gevorg Grigoryan, Amy E. Keating

## Abstract

Understanding the relationship between protein sequence and structure well enough to rationally design novel proteins or protein complexes is a longstanding goal in protein science. The Protein Data Bank (PDB) is a key resource for defining sequence-structure relationships that has supported the development of critical resources such as rotamer libraries and backbone torsional statistics that quantify the probabilities of protein sequences adopting different structures. Here, we show that well-defined, non-contiguous structural motifs (TERMs) in the PDB can also provide rich information useful for protein-peptide interaction prediction and design. Specifically, we show that it is possible to rapidly predict the binding energies of peptides to Bcl-2 family proteins as accurately as can be done with widely used structure-based tools, without explicit atomistic modeling. One benefit of a TERM-based approach is that prediction performance is less sensitive to the details of the input structure than are methods that evaluate energies using precise atomic coordinates. We show that protein design using TERM energies (dTERMen) can generate highly novel and diverse peptides to target anti-apoptotic proteins Bfl-1 and Mcl-1. 15 of 17 peptides designed using dTERMen bound tightly to their intended targets, and these peptides have just 15 - 38% sequence identity to any known native Bcl-2 family protein ligand. High-resolution structures of four designed peptides bound to their targets provided opportunities to analyze strengths and limitations of this approach. Dramatic success designing peptides using dTERMen, which comprised going from input structure to experimental validation of high-affinity binders in approximately one month, provides strong motivation for further developing TERM-based approaches to design.

## Introduction

Protein-protein interactions (PPIs) are central to nearly all biological processes and contribute to pathology in countless human diseases [1]. Reagents that can disrupt PPIs are highly sought for basic research and for therapeutic development, but the size and complexity of many protein interfaces make them difficult to target. For example, large binding sites that have multiple, widely spaced hotspots are notoriously difficult to disrupt with small molecules, as are flat interfaces that lack pockets [2]. Antibodies and nanobodies can block PPIs and have the advantage, relative to small molecules, of binding to larger protein interfaces. The difficulty of delivering large molecules into the cell, coupled with the low stability of some antibody-derived agents in the reducing environment of the cytoplasm, has largely limited their application to extracellular targets or chemically permeabilized cells *ex vivo* [3]. Furthermore, there are PPI interfaces for which antibodies are non-ideal due to the architecture of the immunoglobulin domain [4].

Peptides provide a complementary and highly promising approach to targeting PPI interfaces. Peptide-protein interactions are ubiquitous in nature, where there are many examples of short segments binding to large, structurally complex protein surfaces [5–7]. Peptides can be delivered into cells by chemically modifying them to increase hydrophobicity and hide hydrogen bonds/negative charges [8–10], conjugating them to transduction domains (such as cell-penetrating peptides) [11–13], or delivering them using nanoparticles [14]. Nevertheless, there are obstacles to developing useful peptide inhibitors. Peptides derived from naturally occurring sequences have non-optimal pharmacological properties, because they weren’t evolved for function as reagents or therapeutics. Furthermore, native ligands often have a binding affinity or specificity profile different from what is desired for a given application. Significant sequence optimization is typically required to minimize off-target binding, decrease protease sensitivity, reduce immunogenicity, and improve pharmacokinetics. Because we lack the ability to predict pharmacological potential *a priori*, an ability to rapidly generate numerous diverse peptide sequences that tightly bind/inhibit a target PPI would be transformative for the development of peptide therapeutics.

Current approaches for discovering diverse peptide PPI inhibitors have limitations. State of the art methods rely heavily on experimental screening, and generating a peptide library requires selecting a parent sequence in advance (except for very short peptides). The parent is most often a naturally occurring ligand, around which only a vanishingly small fraction of the sequence space can be queried. Screening that selects for the “best” binders in a population does not typically provide diverse leads. Rational design, e.g. using computational models to search sequence-structure space on a much larger scale, can effectively guide screens to sequences unrelated to those represented in nature. However, given the essentially infinite space to explore and the difficulty of accurately predicting the best binders, the success rates of rational, structure-based methods have been low [15–19].

Recent methodological developments have shown that mining sequence-structure relationships from the Protein Data Bank (PDB) has the potential to improve the efficiency and efficacy of structure-based modeling and design [20–24]. It has long been recognized that proteins are composed of recurring structural elements [25,26]. The large number of solved structures now makes it possible to compile a finite, yet near-complete, list of the recurring tertiary structural motifs (here called TERMs) that are needed to construct any protein structure [27]. Recent analyses have demonstrated that TERMs have characteristic sequence preferences that can be detected by statistical analysis of solved structures [28]. These observations provide the foundation for a formalism that can quantify the compatibility of any sequence with any specified structural scaffold, as described by Zhou *et al*. [29].

TERM-based computational analyses have already demonstrated utility for challenging modeling tasks. For example, a statistical analysis of TERM sequences is remarkably effective at discriminating between good and poor structure predictions, on par with or exceeding leading model quality assessment metrics [30]. Zheng et al. also showed that TERM sequence statistics capture aspects of protein thermodynamics and can be used to predict stability changes upon mutation as well as, or better than, state-of-the-art physics-based or statistical methods [28]. Finally, TERM-based sequence-structure relationships can be applied to protein design. Mackenzie *et al*. showed that choosing optimal sequences for native backbones, based solely on statistics of constituent TERMs, leads to native-like sequences and rationalizes observed evolutionary variation [27]. More recently, Zhou *et al*. described and extensively benchmarked a TERM-based design method, called dTERMen (<u>d</u>esign with <u>TERM</u> <u>en</u>ergies), demonstrating that it is predictive with respect to available data and can generate novel sequences that fold to the intended structure [29]. So far, TERM-based methods have not been applied to predicting or designing protein interactions.

dTERMen is distinct from many other approaches to protein design because it chooses sequences for a target structure based on mining the PDB for TERM-based sequence statistics. These statistics quantify sequence-structure compatibility in the context of ensembles of structurally similar TERMs, as opposed to a single fixed backbone. This approach implicitly accounts for some backbone flexibility, which is advantageous. However, building a scoring function from an ensemble of structures also means that design results are not always easy to interpret in the context of a single ground-state structure. For example, steric clashes that are apparent when a designed sequence is modeled in the context of a fixed backbone structure may or may not be destabilizing.

In this work, we tested the ability of dTERMen to analyze and re-design peptide binders of the important anti-apoptotic proteins Bfl-1 and Mcl-1. Along with paralogs Bcl-2, Bcl-x_L_, and Bcl-w, these proteins promote cellular survival by binding and sequestering pro-apoptotic proteins. Mcl-1 and Bfl-1 have established roles in cancer cell survival and the development of chemoresistance [31,32]. Although blocking Bcl-2 protein binding to pro-apoptotic partners is a validated clinical strategy [33,34], there are no clinically approved inhibitors of Bfl-1 or Mcl-1 at this time. Small molecules, peptides, and mini-proteins have been described as potential inhibitor leads [8,35–38]. For comparison with prior design strategies that required extensive library screening, we tested the ability of dTERMen to generate peptide binders of Bcl-2 family proteins. Our success validates dTERMen as a promising and novel approach for rapid early stage discovery of diverse and high-affinity peptide ligands.

## Results

Bcl-2 family proteins Bcl-2, Bcl-x_L_, Bcl-w, Bfl-1 and Mcl-1 bind to Bcl-2 homology 3 (BH3) motifs within their interaction partners. The short ~23-residue BH3 motif, typically disordered in solution, folds into an alpha helix upon binding. Below, we refer to positions in BH3 peptides using a heptad notation, defined in Table S1 of native BH3 sequences, that reflects the periodicity of the amphipathic helix. In this notation, positions 2d, 3a, 3d and 4a are typically hydrophobic, position 3a is conserved as leucine in native BH3 motifs, position 3e is conserved as a small amino acid, and position 3f is conserved as aspartate.

### Benchmarking interaction prediction performance

To evaluate the potential of dTERMen for designing peptide ligands for Bcl-2 family targets, we tested its performance on a variety of prediction tasks. We used a dataset consisting of 4488, 4648 and 3948 measurements of BH3 peptides binding to Bcl-x_L_, Mcl-1 and Bfl-1, respectively [39]. The peptides were 23 residues in length and contained between 1 and 8 mutations made in the background of the BH3 sequences of human pro-apoptotic proteins BIM or PUMA. Affinity values were obtained using amped SORTCERY, a high-throughput method for quantifying dissociation constants of peptides displayed on the surface of *Saccharomyces cerevisiae* [39,40]. Using this assay, thousands of peptides were determined to have apparent cell-surface dissociation constants ranging from 0.1 to 320 nM, with some peptides classified simply as binding tighter or weaker than the extremes of this range.

Using the amped SORTCERY data, we defined three tasks of increasing difficulty. The easiest task was to discriminate the tightest 20% of binders from the weakest 20%, for a particular target protein. We also defined an enrichment task, which involved identifying the tightest 10% of binders and, finally, the difficult task of predicting quantitative affinities within a 5 kcal/mol range in apparent binding energies. For these tests, we compared the performance of dTERMen with that of commonly used methods Rosetta [41,42] and FoldX [43].

As input for the prediction calculations, we used experimental structures of Bcl-2 protein-peptide complexes. Querying the PDB and filtering for bound peptides of at least 20 amino acids in length yielded 15, 6 and 25 protein-peptide complexes for Bcl-x_L_, Bfl-1 and Mcl-1, respectively (Table S2). An analysis of the BH3 peptides in these complexes revealed that they all adopt a similar binding mode (Table S3) (average pairwise Cα RMSD of 1.64 ± 0.85 Å for the binding interface, defined as Cα atoms of the peptide and surrounding protein residues; see Methods for details).

We first tested whether different modeling approaches could discriminate high affinity binders from peptides that were not observed to bind or that bound weakly. Table 1 reports the average performance of each method over all structural templates. Binary classification of tight binders vs. weak binders is reported as the area under the receiver operating characteristic curve (AUC). An AUC value of 1 corresponds to perfect discrimination and an AUC value of 0.5 corresponds to random guessing. Performance averaged for all protein targets shows that dTERMen (AUC_avg_ = 0.78) has similar predictive power to the other scoring methods, Rosetta (AUC_avg_ = 0.75) and FoldX (R_avg_ = 0.75). The small difference in results is driven by performance on the Bcl-x_L_ dataset, for which dTERMen (AUC_avg_ = 0.75) is better than Rosetta (AUC_avg_ = 0.69) and FoldX (AUC_avg_ = 0.68). For the task of predicting quantitative binding energies, performance averaged for all protein targets shows that dTERMen (R_avg_ = 0.37), Rosetta (R_avg_ = 0.34) and FoldX (R_avg_ = 0.31) gave similar performance, with dTERMen outperforming the other methods for Bcl-x_L_.

**Table 1.** –Interaction prediction performance averaged over all templates ^a^.

|  | AUC<sup>b</sup> |  |  |  | Affinity Correlation<sup>c</sup> |  |  |  | Enrichment of the top 10% of binders (%)<sup>d</sup> |  |  |  |
| --- | --- | --- | --- | --- | --- | --- | --- | --- | --- | --- | --- | --- |
| Target | Bcl-x<sub>L</sub> | Bfl-1 | Mcl-1 | Mean<sup>e</sup> | Bcl-x<sub>L</sub> | Bfl-1 | Mcl-1 | Mean<sup>e</sup> | Bcl-x<sub>L</sub> | Bfl-1 | Mcl-1 | Mean<sup>e</sup> |
| **FoldX** | 0.68<br>±<br>0.11<br>(3)<sup>f</sup> | 0.75<br>±<br>0.06<br>(2) | 0.82<br>±<br>0.08<br>(6) | 0.75 ±<br>0.06<br>(14) | 0.23<br>±<br>0.11<br>(1) | 0.34<br>±<br>0.07<br>(2) | 0.37<br>±<br>0.11<br>(3) | 0.31 ±<br>0.06<br>(9) | 20.3<br>± 6.6<br>(1) | 19.9<br>± 4.0<br>(0) | 23.4<br>± 8.2<br>(0) | 21.4<br>± 1.8<br>(1) |
| **Rosetta** | 0.69±<br>0.05<br>(1) | 0.72<br>±<br>0.02<br>(2) | 0.85<br>±<br>0.03<br>(12) | 0.75 ±<br>0.07<br>(15) | 0.24<br>±<br>0.06<br>(1) | 0.32<br>±<br>0.03<br>(2) | 0.45<br>±<br>0.04<br>(16) | 0.34 ±<br>0.09<br>(19) | 20.5<br>± 3.8<br>(1) | 18.96<br>± 3.6<br>(0) | 24.44<br>± 4.4<br>(13) | 24.3<br>± 6.7<br>(14) |
| **dTERMen** | 0.75<br>±<br>0.06<br>(11) | 0.73<br>±<br>0.03<br>(2) | 0.83<br>±<br>0.06<br>(7) | 0.78 ±<br>0.04<br>(20) | 0.35<br>±<br>0.07<br>(13) | 0.32<br>±<br>0.04<br>(2) | 0.44<br>±<br>0.10<br>(6) | 0.37 ±<br>0.05<br>(21) | 29.9<br>± 4.9<br>(13) | 27.5<br>± 1.4<br>(6) | 35.2<br>±<br>35.1<br>(12) | 30.9<br>± 3.2<br>(34) |

a: Average and standard deviation of the reported performance metric over all templates.
b: Area under the ROC curve for discriminating the top 20% of binders from the bottom 20%.
c: Pearson correlation between predicted and experimental binding energy.
d: Percentage of top 10% of binders found in the predicted top 10% of binders.
e: Average and standard deviation of performance over all three proteins.
f: The number in parentheses is the number of templates for which the indicated method gave the best performance, out of the three methods tested. The total number of templates was 15 for Bcl-x<sub>L</sub>, 6 for Bfl-1 and 25 for Mcl-1.

Many applications seek the tightest binding partners for a protein target, given that these may have the greatest potential as reagents or therapeutics. We used an enrichment test to evaluate methods for their ability to recognize high-affinity binders. Specifically, we used each method to rank the 4386, 4491 or 3805 sequences that had measured affinities for Bcl-x_L_, Mcl-1 or Bfl-1. We then examined the top 10% of computationally ranked sequences to determine what proportion of the top 10% of experimental binders were captured. Overall, dTERMen had better enrichment performance than Rosetta and FoldX (31% vs. 24% and 21%) and performed better than the other methods on most problems. We tested predictions using 45 different input structures, and dTERMen had the best enrichment performance for 34 of these cases. Notably, dTERMen had the best enrichment score on all of the Bfl-1 templates and 13 of the 15 Bcl-x_L_ templates.

The results in Table S2 show that predictive power varies significantly as a function of the template used for modeling. For example, FoldX predictions for the Bcl-x_L_ dataset resulted in AUC values from 0.39 to 0.82, depending on template choice. In one case, multiple protein complexes in the asymmetric unit of one crystal structure (5C6H), with an average pairwise binding interface RMSD of 0.69 Å, gave AUC values from 0.65 to 0.82. It is not surprising that binding affinity predictions depend on the input template structures, particularly for dTERMen and FoldX, which do not perform explicit template structure relaxation. But there is no reliable way to know, a priori, which template will give the best agreement with experiments. One approach could be to choose the crystal structure that has the best resolution. However, we found no relationship between structure quality and performance for any of the methods (Fig. S1). A computationally tractable approach to template selection is to use the template that results in the lowest predicted energy for each sequence. Table 2 shows that, without exception, performance improved for all methods when the lowest energy for each sequence, over all templates, was used. FoldX performance improved the most; FoldX mean AUC increased from 0.75 to 0.85, the mean Pearson correlation for binding affinity values improved from 0.31 to 0.47, and mean enrichment of top binders increased from 21% to 29%.

**Table 2.** –Interaction prediction performance using the template that gave the lowest energy ^a^.

| Target | AUC (%) |  |  |  | Affinity Correlation |  |  |  | Enrichment of the top 10% of binders (%) |  |  |  |
| --- | --- | --- | --- | --- | --- | --- | --- | --- | --- | --- | --- | --- |
|  | Bcl-x<sub>L</sub> | Bfl-1 | Mcl-1 | Mean | Bcl-x<sub>L</sub> | Bfl-1 | Mcl-1 | Mean | Bcl-x<sub>L</sub> | Bfl-1 | Mcl-1 | Mean |
| **FoldX** | 0.83 | 0.83 | 0.93 | 0.85 ± 0.05 | 0.39 | 0.46 | 0.56 | 0.47 ± 0.07 | 27.4 | 27.3 | 32.7 | 29.1 ± 2.5 |
| **Rosetta** | 0.70 | 0.76 | 0.92 | 0.78 ± 0.09 | 0.25 | 0.33 | 0.53 | 0.37 ± 0.12 | 20.1 | 20.0 | 38.1 | 26.0 ± 8.5 |
| **dTERMen** | 0.77 | 0.74 | 0.89 | 0.80 ± 0.06 | 0.38 | 0.33 | 0.50 | 0.41 ± 0.07 | 35.3 | 29.2 | 43.3 | 35.6 ± 5.8 |

a: Notes for Table 1 apply

We were struck by the strong dependence of predicted binding affinities on the choice of template structure and thought this might be an area where dTERMen could provide a modeling advantage, particularly for complexes for which only one or a few structures have been solved. The robustness of prediction performance to very small differences in input structures was evaluated using 294 pairs of closely related structures of the same protein complex that had binding interface C_α_ atom RMSD < 1 Å. For each pair, we computed the correlation of predicted binding energies for all peptides with measured dissociation constants. The results are shown in Fig. 1. On average, dTERMen (R_avg_ = 0.77) is much less sensitive to small differences in input template than FoldX (R_avg_ = 0.55). When run with default options, the Rosetta “relax” protocol is slightly more robust than FoldX (Rosetta R_avg_ = 0.60), although further structural sampling could, at least in theory, lead to a convergence of the Rosetta predictions made using different templates, albeit at a higher cost in computing time.

**Figure 1.**
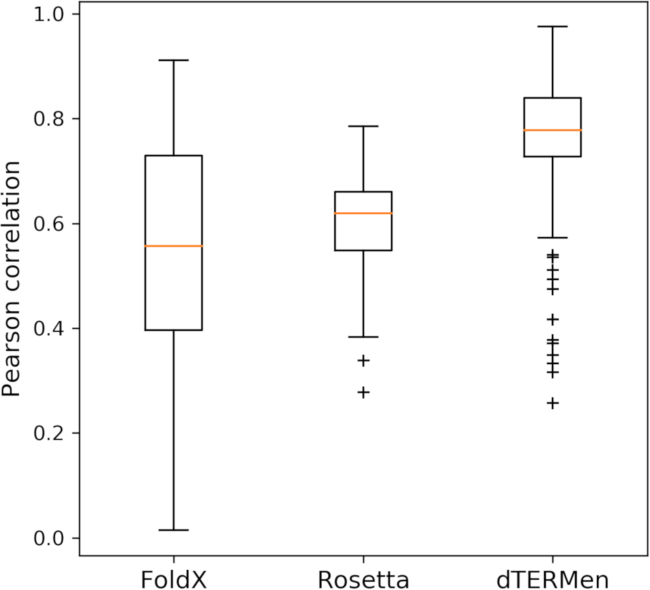
Prediction robustness to small differences in input structures. Distribution of Pearson correlation coefficients (R) comparing scores when modeling on templates with binding mode RMSD within 1 Å. 294 pairs of structures and 6155 unique amped SORTCERY sequences were used to generate the data. The median R value is indicated in red, data between the first and third quartiles (Q1-Q3) are within the box, the whiskers represent extensions of Q1 and Q3 by 1.5 times the interquartile distance (Q1-Q3), and outliers show points outside the whiskers. The larger correlation coefficients for dTERMen results, compared to other methods, indicate lower sensitivity to small structural differences in templates.

### dTERMen designs

Because dTERMen performed at least as well as established scoring functions in benchmarking, we reasoned that it might be useful for designing peptide binders. Given a template structure, dTERMen can be used to solve for the optimal sequence to fit on the template, or in this case to fit on the peptide chain in the template given a fixed sequence for the protein target. We chose 5 structures as design templates: two structures of Bfl-1 complexes and three structures of Mcl-1 complexes (Table S4). Templates were chosen to sample structural diversity, because we observed that designing on different templates provides access to different sequences (Fig. S2).

For Bfl-1-targeted designs, we selected structures of Bfl-1 bound to the natural ligand PUMA (PDB ID 5UUL) and of Bfl-1 bound to a Bfl-1 selective peptide (FS2) that was identified in a previously reported screen (PDB ID 5UUK) [36]. Because the backbones of peptides PUMA and FS2 are shifted 1.2 Å and rotated 17° relative to one another in the Bfl-1 binding pocket, we expected to see differences in the optimal sequences identified by dTERMen for these two templates. For the Mcl-1 targeted designs, we used structures of Mcl-1 bound to the natural ligand BIM (PDB ID 2PQK) and to a chemically crosslinked variant of the natural ligand BID, called BID-MM (PDB ID 5C3F) [44,45]. These two binding modes are similar (peptide RMSD = 0.76 Å when superimposing the binding interface), but the Mcl-1 protein has differences in the binding pocket in the two structures (binding site RMSD = 1.13 Å). We also used a structure of peptide FS2 bound to Mcl-1. FS2 has low affinity for Mcl-1 (K_d_ > 3 μM) but engages the protein in a unique binding pose (PDB ID 5UUM) [36].

Peptide sequences were designed on each of the templates 5UUL, 5UUK, 2PQK, and 5C3F using dTERMen. Preliminary calculations showed that the designed sequences with the best dTERMen scores included medium sized hydrophobic residues at 3a and negatively charged residues at 3f, similar to the conserved leucine and aspartate residues in native BH3 motifs. However, dTERMen-design sequences did not preserve native trends at position 4b. Specifically, the 4b position of many native BH3 peptides is often asparagine, aspartate or histidine, which can serve as an N-terminal helix cap for helix 5 of Mcl-1 or Bfl-1. We noticed that dTERMen chose a variety of amino acids at this position (Lys, Glu, Ser, Ala, Val, Tyr, and Thr). To explore the reason behind this departure from the sequence patterns of native BH3 domains, we extracted the N-terminal helix-capping motif (i.e., N-terminus of helix 5 and the BH3 capping fragment; see Fig. S3) from each template and recovered closely-matching backbone geometries from the PDB. To our surprise, whereas matches made to Bcl-2 family proteins indeed exhibited a strong preference for asparagine or aspartate at the capping position, the frequency of capping residues across other matches were considerably lower (e.g., on average, 6% and 10% for asparagine or aspartate, respectively, in the top ~600 non-Bcl-2 homologous matches). It is therefore not surprising that the apparently strong capping effect in native BH3 helices was not recapitulated in dTERMen designs. While it was unclear whether a capping residue at position 4b would be required or not, we chose to fix this position to either asparagine or aspartate (based on the residue in the design template). BH3 residue 3b can also make a helix-capping interaction. In this case, we imposed the wild-type amino-acid identity in half of the designs (dF1-dF4, dM1-dM4), while allowing this position to vary in the other half (dF5-dF8, dM5-dM7). Two sequences were designed on 5UUM (one for each dTERMen version), without any sequence constraints.

Table 3 shows the optimal designed peptide sequence (the provably best-scoring sequence, given the constraints) for each template structure. For many of the designs, re-packing the protein and peptide sidechains on the rigid-backbone design template using Rosetta showed evidence for predicted steric clashes of varying severity. We used PyMol to visualize regions of possible over-packing, as shown in Figs. S4-S7. Because some backbone relaxation is expected when designing new protein complexes, and because the dTERMen scoring function predicted that the designed sequences are compatible with structures closely related to the design templates, we did not filter the designs using any kind of clash criterion.

**Table 3.**
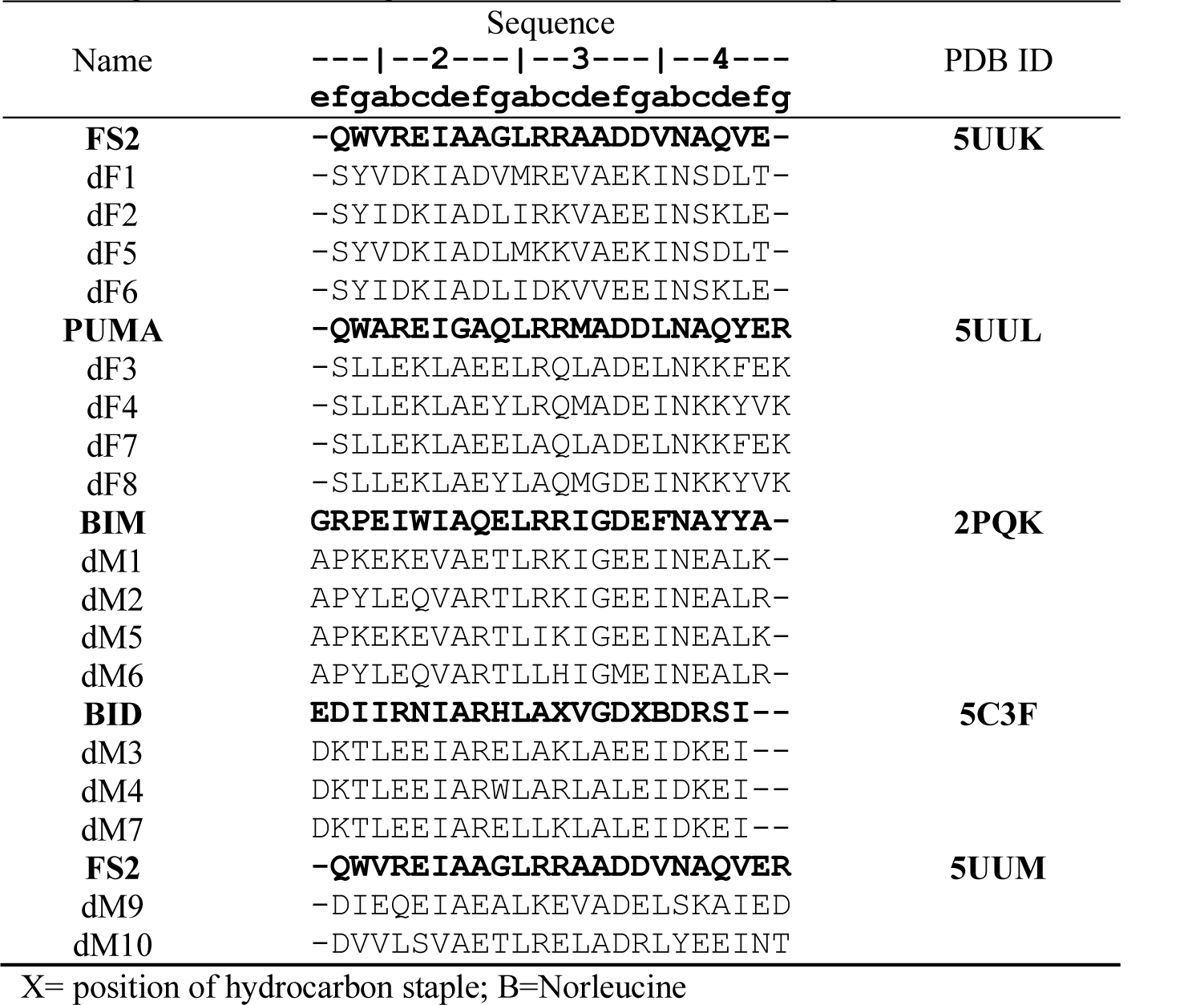
BH3 sequences for template structures (bold) and for peptides designed on those templates using dTERMen. Designs dF1-dF9 target Bfl-1 and dM1-dM10 target Mcl-1.

Fig. 2A shows sequence logos built from 1000 sequences designed on each template, generated from a Monte Carlo simulation without any constraint on position 3b (see Methods). These data, and the designed sequences in Table 3, confirm that peptides designed on different templates were highly distinct, as anticipated. Particularly notable was the diversity observed at positions 3a and 3f. Although dTERMen overwhelmingly chose leucine at position 3a for peptides designed on template 5C3F, matching the conservation observed in native BH3 sequences, greater sequence variation was observed at this site in designs based on other templates. For example, designs based on structure 5UUK included isoleucine or methionine more often than leucine. Position 3f is conserved as aspartate in the natural sequences, but dTERMen chose a variety of polar residues at this site for all templates.

**Figure 2.**
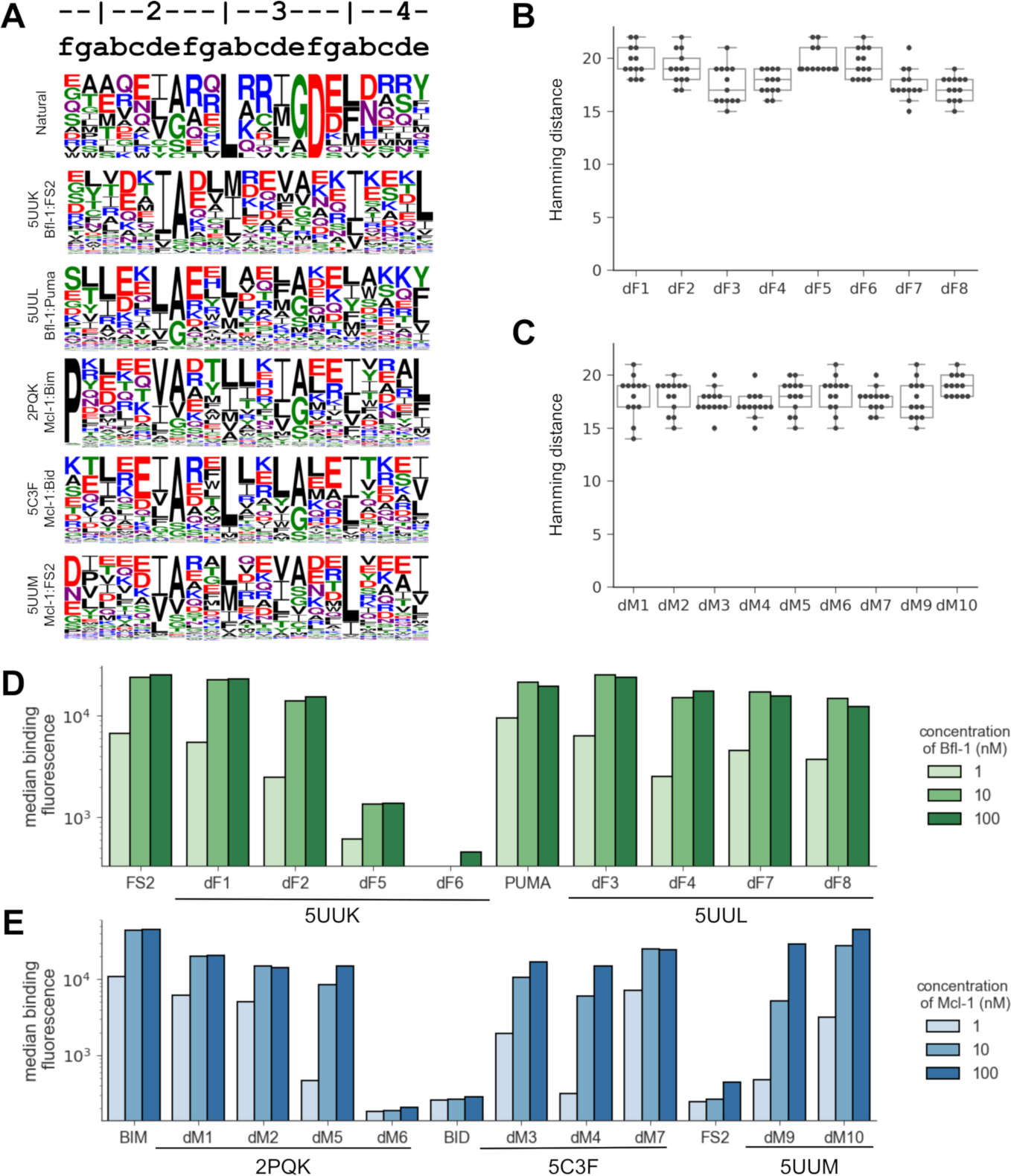
dTERMen design of peptides to bind Mcl-1 and Bfl-1. A) Sequence logos for peptides designed using dTERMen on each of the design templates 5UUK, 5UUL, 2PQK, 5C3F, and 5UUM. Heptad notation for the peptide sequences is shown above the logos. A list of the 13 BH3 motif sequences used to generate the “Natural” logo is in Table S1. B-C) The sequences of the Bfl-1 designs (B) and the Mcl-1 designs (C) were compared to known natural BH3 motifs in Table S1. D-E) The designed sequences were cloned into yeast for cell surface display, and binding to each protein was measured using FACS. Shown here is the median fluorescence binding signal of each peptide in the presence of 1, 10, or 100 nM of the target proteins Bfl-1 (D) or Mcl-1 (E). Data for replicate and off-target measurements are provided in Table S7 and Fig. S8.

To evaluate the predictions made by dTERMen, 17 out of the 18 designed peptides in Table 3 were selected for experimental testing. Sequence dM8, designed on template 5C3F, was not tested because it was only one mutation away from design dM7. The sequences chosen for testing, like all sequences resulting from the design protocol, were very different from any previously known BH3 sequences. Fig. 2B summarizes the minimum number of mutations between the peptides we tested and any of the 13 native BH3 sequences in Table S1 (minimal Hamming distance). Designed peptide binding to Bfl-1, Mcl-1, Bcl-x_L_, Bcl-w, and Bcl-2 was assayed by yeast-surface display. Binding data from yeast-surface display assays have been shown to correlate well with solution affinity measurements, and many BH3 peptides that are tight binders on the yeast cell surface have also been validated as high-affinity binders in solution [35,46,47]. 7 out of 8 peptides designed to bind Bfl-1 showed concentration-dependent binding that saturated at or below 10 nM Bfl-1. 8 of 9 sequences designed to bind Mcl-1 also showed concentration-dependent binding, with apparent cell-surface dissociation constants estimated as < 100 nM (Figs. 2D, E). The results show that constraints on the helix-capping residues at positions 3b and 4b were not necessary for the designed peptides to bind their targets tightly. Peptides designed based on the 5UUM template, a structure of Mcl-1 bound to low-affinity ligand FS2, bound approximately 100-fold more tightly than did FS2 itself, supporting dTERMen as a way to improve the affinity of initial leads for which structures are available (Table S7, Fig. S8).

Peptides dF6 and dM6 did not bind to their targets with high affinity. Peptide dF6 has a valine at position 3e, which is conserved as small (Ala or Gly) in native BH3 peptides, in previously reported designed peptides, and in all of the other dTERMen-designed peptides that we tested [47–49]. Structural matches identified by dTERMen as part of the design process suggested that valine could be accommodated in the context of helix-helix interface geometries highly similar to the one in 5UUK between Bfl-1 residue 88 and BH3 position 3e. In fact, the second-closest match to this local interfacial geometry in our database (backbone RMSD of only 0.27 Å) harbors a valine (Fig. S9). Nevertheless, an all-atom model built using template 5UUK highlights clashes due to the close proximity of the C_μ_ atom of dF6 position 3e and the backbone of position 88 in helix 5 of Bfl-1 (Fig. S4A), and valine may be too large to be accommodated at this site. For design dM6, we hypothesize that substitution of arginine and aspartate at positions 3b and 3f of BIM with leucine and methionine, respectively, and concomitant disruption of a charged network between the peptide and the protein, was destabilizing. These features are consistent with dM6 not binding to any of the Bcl-2 family members we tested (see below).

There is substantial interest in developing Bcl-2 family paralog selective inhibitors [8,31,35,49]. To determine whether our designs cross-react with other anti-apoptotic family members, we tested binding of each peptide to Bfl-1, Mcl-1, Bcl-xL, Bcl-w, and Bcl-2. Interestingly, the Bfl-1 binders that were designed on the structure of PUMA in complex with Bfl-1 (5UUL) bound to multiple Bcl-2 family members. In contrast, peptides designed on 5UUK, which is the structure of Bfl-1-specific peptide FS2 bound to Bfl-1, were > 100-fold selective for Bfl-1, like FS2 itself. The data were less clear for Mcl-1 binders, some of which were selective (dM1, dM5) whereas others were not (dM2, dM3, dM4, dM7, dM9, dM10) (Table S7, Fig. S8).

To determine whether the designed peptides maintained the binding mode of the templates they were designed on, we solved crystal structures for four of the peptides that bound tightly to their targets: dF1 and dF4 in complex with Bfl-1, and dM1 and dM7 in complex with Mcl-1 (Fig. 3). Statistics for data collection and refinement are reported in Table S5. The structure of dF1 in complex with Bfl-1, resolved to 1.58 Å, shows that this peptide binds very similarly to FS2 in template 5UUK (Fig. 3A). It is striking how similar the pocket-facing positions of the designed peptide dF1 and template peptide FS2 are, even though the sequence identity of these two peptides is low (27%) and no information about the FS2 sequence was used in the design process (Fig. S10).

**Figure 3.**
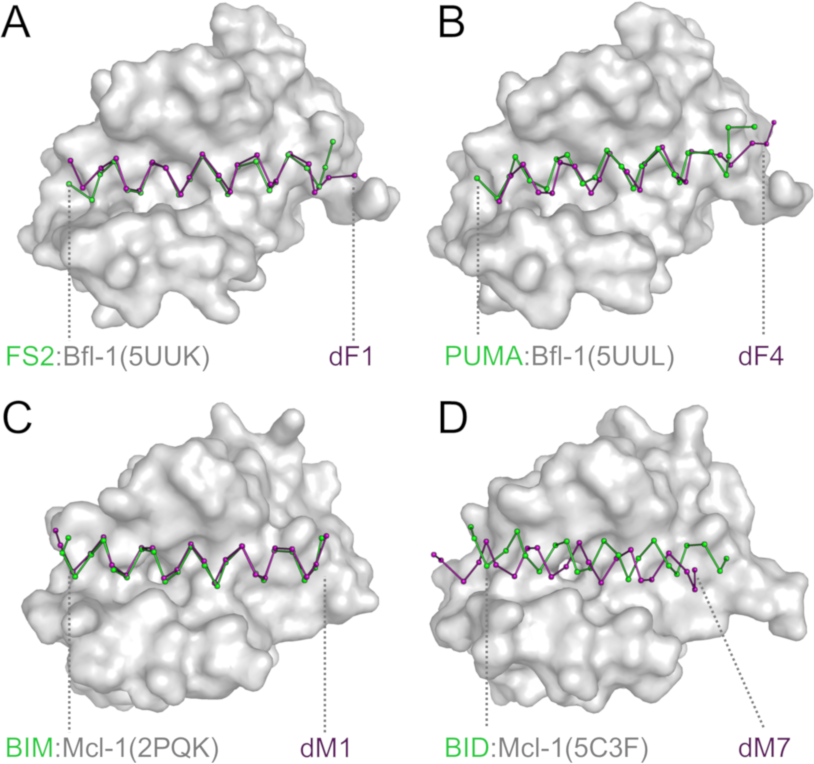
Comparison of the structures of designed complexes and their templates. X-ray crystal structures of (A) dF1 bound to Bfl-1, (B) dF4 bound to Bfl-1, (C) dM1 bound to Mcl-1, and (D) dM7 bound to Mcl-1 (all with the peptide in purple) are compared to the template structures on which they were designed (green ribbon and gray surface). The N-terminal end of each peptides lies to the left in the figure. Modeling dF1 onto the FS2 backbone in structure 5UUK indicated minor clashes, including between methionine at position 3a and residues in the P2 pocket of Bfl-1 (Met 75, Phe 95, and Glu 78), isoleucine at position 4a with Val 44 in helix 2 of Bfl-1, and valine at position 3d with Val 48 and Val 44 of helix 2 of Bfl-1 (Fig. S4C). A more substantial clash was anticipated between valine at position 2g and Leu 52 of helix 2 of Bfl-1 (Fig. S4F). The crystal structure of dF1 bound to Bfl-1 shows how small adjustments accommodate these residues. For example, in the region around valine at 2g, small backbone adjustments are seen for Bfl-1 residues 50-63 that make room for this residue and lead to a modest divergence of the N-terminus of FS2 in 5UUK compared to dF1 in our new structure (Fig. 3A).

We solved the structure of dF4 bound to Bfl-1 to 1.75 Å and found that the C-terminal end of the peptide adopts a different conformation than does PUMA BH3 bound to Bfl-1 in structure 5UUL (Fig. 3B). In template 5UUL, the helix begins to unwind around position 4d, but in the redesigned structure the helix unwinds 3 residues earlier. dTERMen identified relatively few matches for structural elements at the C-terminus of 5UUL, which may have contributed to the deviation from the design template (Fig. S11). At the N-terminus, the sequence of dF4 is very different from that of PUMA; there is only one identical residue within the first 10 residues. An important change was glycine (in PUMA) to alanine (in dF4) at position 2e. In 5UUL, this site is located at a tightly packed helix-helix crossing. Although only glycine can fit when modeled on the rigid design template, TERM statistics indicated that alanine is common in very similar geometries. The solved structure shows how the dF4 helix shifts slightly to accommodate alanine, along with other sequence changes.

We solved the structure of dM1 bound to Mcl-1 to 1.95 Å and found that that it bound very similarly to the BIM BH3 peptide in design template 2PQK (Fig. 3C). However, the structure of dM7 bound to Mcl-1 at 2.25 Å resolution revealed a substantial change in the binding mode of the peptide (Fig. 3D, Fig. S12A). The helix is shifted in the groove by 3.43 Å and rotated by 19 degrees along the helix axis, relative to the position of BID-MM in the design template structure 5C3F. A shift of the helix in the groove by approximately one-half helical turn re-positions leucine at 3a relative to what is observed in structures of native BH3 peptides bound to Bcl-2 family proteins. Furthermore, the canonical BH3 interaction of aspartate at 3f with Bfl-1 Arg 263 is replaced by a salt bridge with an aspartate at position 4b in the peptide (Fig. S12B). In Mcl-1, alpha helix 4 is rearranged relative to its position in the template, to accommodate the unusual sequence. The reorganization may have resulted from introducing two leucine residues at peptide positions 3b and 3f. Not only does leucine at 3f remove the aspartate residue at this position in BIM, BID and PUMA, but leucine at 3b is predicted to interfere with an intra-molecular salt bridge between Bfl-1 residues 256 and 263. The shift of peptide dF1 observed in the crystal structure restores the salt-bridge network between Bfl-1 and the peptide, using a different peptide residue, as shown in Fig. S12B. One complication in evaluating this structure is that there are close crystal-packing contacts between two copies of the Mcl-1:dM7 complex, near the C-terminal end of the binding groove, and involving alpha helix 4 of Mcl-1 (Fig. S13). We cannot rule out the possibility that crystal packing forces favored population of a minor structural species, and that the designed binding mode may be populated in solution.

In summary, x-ray crystallography revealed that backbone positioning of two of the crystalized designs (dF1 and dM1) were sub-Ångstrom matches to their design templates, over most of the length of the peptide. Another peptide (dF4) bound in a geometry that shared high similarity with its template, but the remaining design (dM7) bound in an unexpected, dramatically shifted orientation.

## Discussion

Using dTERMen, we were able to rapidly design high-affinity binders of Bcl-2 family proteins without the need for explicit modeling of complex structures or expensive experimental library screening. Previous work has shown that this is not a trivial task. For example, in a library of random peptides, nearly all fail to bind Mcl-1 detectably [50]. Additionally, even in carefully designed libraries containing peptides with fewer than 6-8 mutations compared to natural BH3 domains, most sequences fail to bind Bfl-1 and Mcl-1 [36]. In contrast, using dTERMen, we found that 15/17 of the designs bound with native-like affinity, even though the sequences were 14-22 mutations away from known BH3 binders (Fig. 2 B-C).

Our design protocol provided access to novel and diverse sequences. Some of the tight binders we discovered using dTERMen lack the highly conserved leucine and aspartate residues common to all known, native BH3 sequences (Table 3, Fig. 2A). Not only do our results suggest that these residues are not necessary for binding, but they show that dTERMen is a useful tool for discovering binders that cannot be predicted based on conserved sequence features. Designing on different structural templates gave rise to different solutions, as illustrated in Fig. 2A. This may seem to be at odds with our finding that dTERMen is robust to small differences in input structure (Fig. 1), but we deliberately chose design templates to sample different peptide docking geometries. We expected these templates to match with different TERMs from the PDB, and thus to generate different sequence predictions. Templates 5UUL and 2PQK are structures of complexes with native, tight BH3 peptide binders (reported dissociation constants of ~1 nM) [36,44]. Other templates we tested, 5C3F and 5UUM, featured peptides that bound their targets more than 3 orders of magnitude more weakly [36,45]. It is notable that template structures for both high-affinity and low-affinity peptide complexes led to novel, high-affinity peptide binders when used as input to dTERMen. Designing on other solved structures could provide access to even greater diversity (Fig. S2). Going beyond solved structures, it may be possible to perform dTERMen design on predicted structures with binding modes that have not been previously observed.

A set of designs with diverse sequences is more valuable that a single design optimized for affinity because it provides opportunities to optimize pharmacological properties not related to binding. Our designed peptides have formal net charges ranging from −7 to +1, predicted helical content ranging from 0.7 to 69.7% and predicted hydrophobicity of 0.03 to 0.48 (Table S5). These properties could affect whether these peptides are disruptive to membranes and how readily they can be delivered to cells. Several studies have shown that the cell permeability of stapled helical peptides depends on peptide properties including charge and hydrophobicity [8,10]. Different sequences will also have different cross-reactivity, immunogenicity, and protease sensitivity, so having many options to choose from increases the chances of developing useful reagents and lead therapeutics. Interestingly, design using dTERMen is compatible with imposing constraints on peptide properties such as net charge, so if the desired physical characteristics of a peptide inhibitor are known, they can be used to direct the search into promising sequence spaces.

The dTERMen scoring potential is based on sequence statistics for structural elements observed repeatedly in nature. There is no formal relationship between these statistics and protein stability or affinity, so the scoring may reflect any number of evolutionary pressures including stability, specificity, folding kinetics, solubility, or other factors. We interpret the success of dTERMen as evidence that whatever evolutionary forces may be contributing to the statistics, there must be a substantial contribution from the free energy of the sequence adopting the evaluated structure. The fact that we designed helix-helix interactions in this project, which are common in the PDB, may be part of the reason dTERMen designs performed so well. Because more structures are deposited in the PDB every day, we expect the range of accessible design targets to increase over time [28].

One attractive feature of dTERMen is that it doesn’t require explicit structural modeling or minimization; the design optimization is performed in sequence space. Although the PDB structure-mining that is required to build the scoring function can be somewhat time consuming (e.g. it takes 7 to 12 CPU hours to generate scoring functions for the structures we analyzed here), once such a function is derived, it is possible to perform design, or to evaluate millions of sequences, in seconds. Another advantage of dTERMen is that there is a structural “fuzziness” built in, because the sequence statistics used for modeling are derived from close, but not exact, matches of TERMs. This makes the method more robust than FoldX to small variations in input structure, as shown in our benchmark testing, and also accounts for some amount of backbone relaxation. In this work, we observed multiple examples where a mutation was accommodated that would not have been designed if modeling was performed on a rigid scaffold (Figs. S4-S7). On the other hand, dTERMen design failures may result from over-packing the protein-peptide interface beyond what can be accommodated by small structural rearrangements. This may be what happened for dF4, the structure of which diverged from the design template structure at the C-terminal end of the peptide, and for dF6, which did not bind tightly to Bfl-1. Future design studies will help calibrate the methods so that diverse sequences can be obtained with reliably high success rates. Combining dTERMen with a post-analysis procedure that includes all-atom modeling with aggressive conformational search, using peptide redocking [51] or MD simulation [52], could be one way to recognize sequences or mutations that can or cannot be accommodated. Although this would increase the computational costs, such a secondary evaluation could be performed for a modest number of promising candidates designs.

One unexpected result from this work is that the specificity profiles of the designs were template dependent. This is particularly striking in the case of design on the FS2 template. Although no off targets were considered during design, the peptides designed using the FS2 structure were highly Bfl-1 selective. In fact, these peptides provide outstanding leads for development as Bfl-1 targeting agents. The specificity of peptides dF1, dF2 and dF5 may be a result of the unique way FS2 engages Bfl-1. FS2 adopts a non-canonical binding mode that has not been observed for natural BH3 ligands [36]. It may be that the interactions with Bfl-1 that support the FS2 binding mode are under less evolutionary pressure to mirror those required for BH3 binding in the other family members, and are thus more likely to be unique (Fig. S14). This is consistent with the idea that a peptide that makes contacts outside of the conserved binding cleft can use these contacts to achieve intra-family specificity [37,53].

This proof-of-principle study makes us enthusiastic about the potential of dTERMen for designing peptide binders and inhibitors. The ease of use, fast run times, and very high success rates on a difficult problem provide compelling evidence of the promise of this approach. Future applications could exploit dTERMen scoring speed by screening proteomes to predict candidate binding partners, or could leverage the robustness of dTERMen to scaffold variation by designing on low resolution structures. There are ample opportunities to improve dTERMen further, for example by combining this sequence-based design approach with all-atom modeling to better assess what mutations can be accommodated by structural relaxation. We look forward to tackling increasingly difficult problems and moving the use of TERM statistics into the mainstream of modern protein design.

## Methods

### dTERMen design scoring function

A full description of the dTERMen procedure, along with extensive validation and benchmarking, is given in Zhou *et al*. [29]. For completeness, we briefly outline the method here, at a high level. Given a target protein structure, *D*, for which an appropriate amino-acid sequence is needed, dTERMen begins by defining effective self energies for each amino acid at each position of *D* and effective pair interaction energies between amino acids at pairs of positions. We collectively refer to these as energy parameters (EPs) and their values in our procedure are deduced from statistics of structural matches to appropriately defined TERMs that make up *D*. The matches are obtained by searching a structural database. In this work, the database was a subset of the PDB containing 14,546 chains from X-ray structures with resolution better than 2.6 Å, pruned for redundancy at 30% sequence identity. Importantly, this means there was no quaternary structural information present in the database, and all insights on how to design domain peptide interfaces were derived from intra-chain examples.

The fundamental idea behind our procedure is to define TERMs from *D* in a way that is targeted at isolating individual EP contributions. For example, to capture the pairwise dependence between amino-acid identities at positions *i* and *j* (i.e., the pair EP), we define a TERM that consists of residues *i*, *j*, and their surrounding backbone fragments (e.g., ± 2 residues around each residue). By obtaining a sufficiently large list of closest matches to the generated motif (pruned for redundancy), one can analyze the co-dependence between identities at *i* and *j*. One complicating factor is that identities at the two positions are also biased by the specific environments from which the matches originate. And, in some cases, this bias could affect the apparent co-dependence. E.g., if the two positions are usually either both buried or both exposed within matches, it may appear that there is a direct favorable interaction between amino acids of similar hydrophobicity at *i* and *j*. Such effects are corrected for in dTERMen by computing EPs as log-odds ratios between observed and expected numbers of observations (e.g., observations of amino-acid pairs in this case), where the expectation is calculated by accounting for the effect of the environment in the structures from which matches originate. Self-EP contributions arising from interactions between a residue and nearby backbone fragments are computed similarly. These include interactions with both the local sequence-contiguous backbone (the own-backbone energy) and backbone fragments proximal in 3D (the near-backbone energy). These contributions augment pre-tabulated amino-acid self-energies associated with different backbone φ/ψ and ω dihedral angles and burial states to form the final EP contributions.

The above computed contributions are compiled into an energy table of one‐ and two-body contributions, after which Integer Linear Programing (ILP) is used to identify the sequence with the most optimal score [54,55]. Note that all energies are defined on the sequence level, such that optimization can proceed directly in sequence space, without the need to build explicit atomic structures. And yet, because each EP contribution arises from an ensemble of TERM matches, a certain amount of implicit backbone flexibility is built into the scoring function.

### dTERMen sequence design protocol

When the design problem pre-specifies some of the residues in the target structure *D*, as is the case in the present application, the calculations remain the same but some re-shuffling between pair and self EPs takes place. For example, when position *i* in an interacting pair *i*-*j* is fixed, the TERM-derived effective pair EP between the two is added to the self-energy of position *j* in the final table. Because in the present case the sequence of the entire domain was always fixed, the only pairwise contributions in the final table were those between pairs of peptide positions.

The two versions of dTERMen used here differ in how TERMs for computing the near-backbone energy for residue *i* are defined (see Zhou *et al*. for full details [29]). The ideal TERM for this purpose would include the residue *i*, its local backbone fragment, all residues with the potential to interact with *i* (through either side-chain or backbone‐‐i.e., influencing residues), and their respective local backbone fragments. If such a TERM has a sufficient number of close structural matches in the database, then this definition works well and the two dTERMen versions will both pick this motif (producing the same result). Because near-backbone TERMs can have many segments (e.g., three potential interacting positions would give rise to a four-segment TERM), they may not always be represented well enough in the database to derive confident sequence statistics on the amino-acid preferences at *i*. In this case, one is forced to consider the effect of the local backbone geometry on position *i* as an aggregate of effects from sub-motifs, and the two versions deal with this differently. Version 35 attempts to identify large sub-motifs, each consisting of *i* and as many of the influencing residues as possible (along with local backbones), such that sub-motifs do not overlap and together cover all influencing residues. This takes a considerable amount of database searching, as many trial submotifs have to be queried. Version 34 speeds this process up, at the cost of some detail, by considering just one sub-motif that includes the most “important” influencing residues (assessed via our geometric measure of contact degree [30]), on the assumption that this motif dominates sequence statistics.

### Structural model generation

We used pyRosetta [56] (Linux release r53335) to generate structural models for dTERMen-designed sequences emergent from ILP optimization. This was done by performing fixed-backbone side-chain repacking of all residues in the domain-peptide complex (peptide residues taken from the dTERMen-optimized sequence) using the talaris2013 forcefield [56] and default parameters in pyRosetta via “standard_packer_task” and “PackRotamersMover” objects. For residues where there was evidence of crowding, all backbone-dependent rotamers of a residue of interest were manually inspected using PyMol. Figs. S4-S7 were made by choosing the least clashing rotamer.

### Sequence logo generation

In addition to obtaining the dTERMen-optimal sequence for each template by ILP, we also performed Monte Carlo (MC) sampling to generate an ensemble of well-scoring sequences as a way of better characterizing the predicted favorable sequence space (see Fig. 2A). To this end, we ran 1000 independent MC trajectories for each template starting with a random sequence. Each trajectory involved 100,000 iterations, at each of which a random mutation was evaluated for acceptance according to the Metropolis criterion. The sampling was performed at constant temperature with kT equal to 1 (this was also the temperature used to derive dTERMen statistical energies). The final accepted sequence from each of the 1000 trajectories was used to build an MSA for each template and to generate the logos in Fig. 2A using WebLogo [57]. No constraint was imposed at position 3b.

### Designed-peptides property prediction

Predicted helical content for designed peptides was obtained from the AGADIR web server [58]. Predicted net charges and hydrophobicity were obtained using the HelixQuest server [59].

### Analysis of similarity of peptide interactions with Bcl-2 family paralogs

The Bfl-1 sequence was aligned with the sequences of Bcl-xL, Mcl-1, Bcl-2, and Bcl-w using ClustalW [60]. Each residue in Bfl-1 was scored for sequence similarity to the corresponding residue in each of the other proteins using the Blosum62 matrix [61]. Substitutions with scores ≥ 0 were considered similar. To display amino-acid conservation at each position on the Bfl-1 structure, as shown in Fig. S14, each residue was colored by the number of proteins with amino acids similar to the one in Bfl-1 at that position.

### Automatic download and annotation of Bcl-2 protein-peptide complex structures

Uniprot sequences for human Bcl-x_L_, Bfl-1 and Mcl-1 were retrieved from Uniprot [62] and blasted against the PDB database [63] (7 Nov 2017). Matched structures were downloaded and standardised by transforming selenomethionine to methionine and removing hydrogens and atoms designated as HETATOM. Sequences were aligned and renumbered based on their corresponding Uniprot template sequence using Needle [64]. Regions that were not matched or that were poorly aligned with the Uniprot sequence were removed from the structure. Chains of length 20-39 residues with more than 30% of their Voronoi surface in contact with the Bcl-2 proteins were identified as interacting peptide [65]. Unless specified, peptides containing non-natural amino acids were removed from the dataset. Only the first model in deposited NMR ensembles was retained. If a structure included multiple complexes in the asymmetric unit, these were split into new files and analyzed separately.

### Alignment on the binding site and method for comparing peptide binding geometry

For every complex, residues within 8 Å of any peptide atom were considered part of the binding interface and all complexes were structurally aligned using only their binding interface C_α_atoms, using 3DCOMB [66]. To automatically define a common reference residue for all bound peptides, we used a graph-based procedure. Each peptide C_α_in the set of superimposed binding interfaces was represented as a node, and an edge was created if the distance between 2 nodes was below a threshold. The distance threshold was initially set at 2 Å and gradually increased by 0.1 Å until the largest clique in the graph included one residue from each complex. This clique represented a set of C_α_ atoms - one in each structure - all within a distance threshold. Residues in this largest clique were arbitrarily given peptide residue number 100; this reference residue corresponds to residue 95 in structure 3FDL. Using this registry, peptides were trimmed to generate a 20-residue long segment chosen by structural inspection to include positions that make extensive contacts with the protein and that are unlikely to be influenced by crystal contacts in the templates used for modeling. This region corresponds to peptide positions 86 to 105 in structure 3FDL. Structures without a complete 20-mer peptide were not used. Binding interfaces were redefined using trimmed peptides, by taking all peptide atoms plus protein residues within 8 Å of any peptide atom.

### Scoring protein-peptide interactions

Structural scoring functions dTERMen (described above), FoldX4.0 and Rosetta were tested for their ability to predict peptide-protein binding affinity using binding data obtained using the SORTCERY protocol [40–43]. Scoring was based on trimmed-peptides structures. Each structure was used as a template input for dTERMen, leading to a scoring function for that template, i.e. a function that can score any peptide binding to the target protein in the template-structure binding mode. FoldX4.0 was used to predict binding affinity by first using FoldX4.0’s “repair” function. Then, for each peptide in the SORTCERY dataset, the repaired template was transformed using the “mutate” function to generate the sequence of the peptide query and scored using the “complex” function. For Rosetta scoring, complex structures generated by FoldX were relaxed with Rosetta (Nov 2017 version rosetta_bin_linux_2017.08.59291, “relax” command) using Talaris2014 or BetaNov force fields [42]. The default parameters of 5 minimization cycles consisting of 4 rounds of repacking were used for the relaxation protocol. Relaxed structures were run through the Rosetta InterfaceAnalyzer module, and the “dG_separated” values were used as the predicted binding energy. This score describes the difference in Rosetta energy of interface residues between the complex structure and corresponding separated chains. For the sake of simplicity in the reporting of benchmarking results, only the latest scoring function of Rosetta (BetaNov) and dTERMen (35) are discussed. dTERMen scoring function 34 and Rosetta Talaris2014 force field yield similar benchmark performance as these newer versions and values can be found Table S2.

### Interaction prediction benchmark

The predictive power of the different structural scoring functions and protocols was assessed by three metrics. First, each method’s ability to discriminate the top 20% tightest-binding peptides from the 20% weakest binders was assessed by calculating Area Under the Curve (AUC) of the Receiver operating characteristic (ROC) curve. Next, precision was evaluated by calculating the correlation between the binding energy determined by SORTCERY, in kcal/mol, and each method’s predicted binding energy (in arbitrary units). Finally, we computed the percentage of the top 10% of binders from SORTCERY experiments that were found in the top 10% of predicted binders. Multiple templates were tested for each protein, and predictive power was evaluated for each template individually. The average performance and standard deviation of performance over all templates was computed and represents the expected value if a random template is chosen. We also report prediction performance using the template that gave the lowest energy for each sequence.

### Protein and peptide purification

Myc-tagged human Mcl-1 (residues 172-327), Bfl-1 (residues 1-151), Bcl-2 (residues 1-217), Bcl-w (residues 1-164), and Bcl-x_L_ (residues 1-209) were used for binding assays. Untagged Bfl-1 (residues 1-151) and Mcl-1 (residues 172-327) were used for crystallography. The proteins used in this study were purified as previously described [47] and frozen at −80 °C. The peptides used for crystallography were synthesized at the MIT biopolymers facility with N-terminal acetylation and C-terminal amidation and were purified by HPLC on a C-18 column with a linear gradient of acetonitrile and water. Purified peptides were lyophilized and resuspended in DMSO. Peptide masses were confirmed by MALDI-TOF mass spectrometric analysis.

### Yeast clones

EBY100 yeast cells were transformed using the Frozen-EZ Yeast Transformation II Kit (Zymo Research) according to the manufacturer’s protocol. For a plasmid backbone, we used the PUMA PCT plasmid [36] and digested it with XhoI (NEB) and NheI-HF (NEB) according to the manufacturer’s protocol. The inserts were constructed by PCR using primers that encoded the peptide sequence flanked with at least 40 bp of the plasmid sequence on either side of the insertion site to facilitate homologous recombination. The inserts and plasmid backbones were mixed at a 5 to 1 ratio for transformation. The transformation mixture was spread onto SD + CAA plates (5 g/L casamino acids, 1.7 g/L yeast nitrogen base, 5 g/L ammonium sulfate, 10.2 g/L Na_2_HPO_4_-7H_2_O and 8.6 g/L NaH_2_PO_4_-H_2_O, 2% glucose, 15-18 g/L agar, 182 g/L sorbitol) and grown at 30 °C for 2 to 3 days. To confirm each strain, colony PCR followed by sequencing was performed on single colonies. Sequence verified colonies were grown overnight in SD + CAA (5 g/L casamino acids, 1.7 g/L yeast nitrogen base, 5 g/L ammonium sulfate, 10.2 g/L Na_2_HPO_4_-7H_2_O and 8.6 g/L NaH_2_PO_4_-H_2_O, 2% glucose). The saturated overnight cultures were diluted with 40% glycerol to a final glycerol concentration of 15% and stored at −80 °C.

### Yeast growth and FACS analysis

A small amount of frozen culture was scraped from the top of frozen culture stocks to inoculate SD + CAA. After passaging overnight at 30 °C, cultures were diluted to an OD600 of 0.005-0.01 in SD + CAA and grown to an OD600 of 0.1– 0.6. Cell cultures were then diluted 25-fold with SG + CAA (5 g/L casamino acids, 1.7 g/L yeast nitrogen base, 5.0 g/L ammonium sulfate, 10.2 g/L Na_2_HPO_4_-7H_2_O and 8.6 g/L NaH_2_PO_4_-H_2_O, 2% galactose) to induce peptide expression and grown for 20-24 hr at 30 °C. To measure binding to surface-displayed peptides, cells were filtered with a 96-well plate filter (10^5^-10^6^ cells/well), washed twice with 150 μL BSS (50 mM Tris pH 8, 100 mM NaCl, 1 mg/ml BSA), and resuspended in BSS with least 10-fold molar excess target protein and incubated in the filter plate for 2 h at room temperature with gentle shaking for equilibration. Binding of the designs to the five Bcl-2 family proteins was measured at 1000 nM, 100 nM, 10 nM, and 1 nM target protein. To detect cell surface expression and binding of target protein, cell suspensions were filtered, washed twice in chilled BSS, resuspended in a 35 μL of 1:100 dilution of primary antibodies (mouse anti-HA, Roche, RRID:AB_514505 and rabbit anti-c-myc antibodies, Sigma, RRID:AB_439680) in BSS and with gentle shaking for 15 min at 4 °C. Cells were then filtered, washed twice in 150 μL chilled BSS, resuspended in 35 μL of a solution of secondary antibodies in BSS (1:40 dilution of APC rat anti-mouse, BD, RRID:AB_398465 and 1:100 dilution of PE goat anti-rabbit, Sigma, RRID:AB_261257) and incubated with gentle shaking in the dark for 15 min at 4 °C. Cells were filtered and washed twice more in 150 μL chilled BSS to remove unbound antibodies. Labeled cells were resuspended in BSS and analysed using a BD FACSCanto with FACSDiva software. The median binding signals of expression-positive cells are shown in Fig. 2D and E, Table S7, and Fig. S8.

### Crystallography

Crystals of Bfl-1 in complex with the designed peptides were grown in hanging drops. To set the drops, untagged Bfl-1 (8 mg/mL in 20 mM Tris, 150 mM NaCl, 1% glycerol, 1 mM DTT, pH 8.0) was mixed in equal molar ratio with the designed peptides. 1.5 μL of the Bfl-1/peptide mixture was pipetted onto a glass coverslip and mixed with with 1.5 μL of well solution (1.8 - 2.0 M NH_4_SO_4_, 50 mM MES pH 6.5). To cryoprotect the crystals, they were transferred into a solution of 2.0 M LiSO_4_ with 10% glycerol. Crystals were flash frozen in liquid nitrogen. Diffraction data were collected at the Advanced Photon Source at the Argonne National Laboratory, NE-CAT beamline 24-ID-C. The datasets were refined to 1.59 Å and 1.75 Å and scaled using HKL2000 [67]. Phenix was used to phase with the Bfl-1 chain from PDB id 5UUK [36,68]. The peptides were modeled into the difference densities using Coot [69]. Iterative rounds of refinement and model building were performed using Phenix and Coot [68,69].

Crystals of Mc1-1 in complex with the designed peptides were grown in hanging drops. To set the drops, TCEP (100 mM) and ZnSO_4_ (50 mM) was added at 10% volume to untagged Mcl-1 (8.5 mg/mL in 20 mM Tris, 150 mM NaCl, 1% glycerol, 1 mM DTT, pH 8.0) before adding equal molar amounts of the designed peptides. To grow crystals of Mcl-1 in complex with dF1, 1.5 μL of the peptide protein mixture was mixed with 1.5 μL of well solution (25% PEG 3350, 50 mM BIS-Tris pH 8.5, 50 mM NH_4_CH_3_CO_2_). Crystals were cryoprotected by adding 3 μL of a solution of 37.5% glucose in 25% PEG 3350, 50 mM BIS-Tris pH 8.5, 50 mM NH_4_CH_3_CO_2_ directly to the drop 0.5 uL at a time. To grow crystals of Mcl-1 in complex with dF7, 2.5 μL of the peptide protein mixture was mixed with 0.5 μL of well solution (1.4 M sodium citrate pH 6.5, 0.1 M HEPES pH 7.5). For cryoprotection, crystals were transferred to 1.6 M sodium citrate pH 6.5, 0.1 M HEPES pH 7.5. Crystals were flash frozen in liquid nitrogen. Diffraction data were collected at the MIT x-ray core facility. The datasets were refined to 1.95 Å and 2.25 Å and scaled using HKL2000 [67]. Phenix was used to phase with the Mcl-1 chain from PDB ID 3PK1 [68,70]. The peptides were modeled into the difference densities using Coot [69]. Iterative rounds of refinement and model building were performed using Phenix and Coot [68,69].

### Availability of scripts and data

For information about how to use dTERMen see grigoryanlab.org/dtermen. Scripts used for the prediction benchmark, protein structure files, predicted energy values, and experimental data can be downloaded from a GitHub repository: https://github.com/KeatingLab/dTERMen_design under the MIT License.

## Acknowledgements

This project was supported by NIGMS award R01 GM110048 to A. Keating and supported by NIH award P20-GM113132 and NSF award DMR1534246 to G. Grigoryan. J. Jenson was partially supported by a fellowship from the Koch Institute for Integrative Cancer Research and V. Frappier was supported by NSERC and FRQNT postdoctoral funding. Part of this work was conducted at the Northeastern Collaborative Access Team beamlines, which are funded by the National Institute of General Medical Sciences from the National Institutes of Health (P30 GM124165). The Pilatus 6M detector on 24-ID-C beam line is funded by a NIH-ORIP HEI grant (S10 RR029205). This research used resources of the Advanced Photon Source, a U.S. Department of Energy (DOE) Office of Science User Facility operated for the DOE Office of Science by Argonne National Laboratory under Contract No. DE-AC02-06CH11357. The content is solely the responsibility of the authors and does not necessarily represent the official views of the National Institutes of Health or the U.S. Department of Energy.

We thank the Koch Institute Flow Cytometry Core Facility for assistance with FACS sorting and the MIT Structural Biology Core Facility, R. Grant for assistance with X-ray crystallography, and members of the Drennan laboratory for help with X-ray data collection. We thank V. Xue for FACS data-processing scripts. We thank the Northeastern Collaborative Access Team beamlines and the Advanced Photon Source, a U.S. Department of Energy (DOE) Office of Science User Facility operated for the DOE Office of Science by Argonne National Laboratory.

## References

1. Chatr-Aryamontri A, Oughtred R, Boucher L, Rust J, Chang C, Kolas NK, et al. The BioGRID interaction database: 2017 update. Nucleic Acids Res. 2017;45: D369–D379. doi:10.1093/nar/gkw1102

2. Arkin MR, Tang Y, Wells JA. Small-molecule inhibitors of protein-protein interactions: progressing toward the reality. Chem Biol. 2014;21: 1102–14. doi:10.1016/j.chembiol.2014.09.001

3. Chames P, Van Regenmortel M, Weiss E, Baty D. Therapeutic antibodies: successes, limitations and hopes for the future. Br J Pharmacol. 2009;157: 220–33. doi:10.1111/j.1476-5381.2009.00190.x

4. Eckert DM, Shi Y, Kim S, Welch BD, Kang E, Poff ES, et al. Characterization of the steric defense of the HIV-1 gp41 N-trimer region. Protein Sci. 2008;17: 2091–100. doi:10.1110/ps.038273.108

5. Kuang X, Dhroso A, Han JG, Shyu CR, Korkin D. DOMMINO 2.0: Integrating structurally resolved protein-, RNA-, and DNA-mediated Macromolecular interactions. Database. 2016;2016: 1–12. doi:10.1093/database/bav114

6. Frappier V, Duran M, Keating AE. PixelDB: Protein-peptide complexes annotated with structural conservation of the peptide binding mode. Protein Sci. 2018;27: 276–285. doi:10.1002/pro.3320

7. Tompa P, Davey NE, Gibson TJ, Babu MM. A million peptide motifs for the molecular biologist. Mol Cell. 2014;55: 161–9. doi:10.1016/j.molcel.2014.05.032

8. Rezaei Araghi R, Bird GH, Ryan JA, Jenson JM, Godes M, Pritz JR, et al. Iterative optimization yields Mcl-1-targeting stapled peptides with selective cytotoxicity to Mcl-1-dependent cancer cells. Proc Natl Acad Sci U S A. 2018;115: E886–E895. doi:10.1073/pnas.1712952115

9. Walensky LD, Bird GH. Hydrocarbon-Stapled Peptides: Principles, Practice, and Progress. J Med Chem. 2014;57: 6275–6288. doi:10.1021/jm40116753481–3495. doi:10.1074/jbc.M116.756718

10. Bird GH, Mazzola E, Opoku-Nsiah K, Lammert MA, Godes M, Neuberg DS, et al. Biophysical determinants for cellular uptake of hydrocarbon stapled peptide helices. Nat Chem Biol. 2016;12: 845–52. doi:10.1038/nchembio.2153

11. Schwarze SR, Ho A, Vocero-Akbani A, Dowdy SF. In vivo protein transduction: delivery of a biologically active protein into the mouse. Science. 1999;285: 1569–72.

12. Nischan N, Herce HD, Natale F, Bohlke N, Budisa N, Cardoso MC, et al. Covalent attachment of cyclic TAT peptides to GFP results in protein delivery into live cells with immediate bioavailability. Angew Chem Int Ed Engl. 2015;54: 1950–3. doi:10.1002/anie.201410006

13. Qian Z, Martyna A, Hard RL, Wang J, Appiah-Kubi G, Coss C, et al. Discovery and Mechanism of Highly Efficient Cyclic Cell-Penetrating Peptides. Biochemistry. 2016;55: 2601–12. doi:10.1021/acs.biochem.6b00226

14. Kumar M, Gupta D, Singh G, Sharma S, Bhat M, Prashant CK, et al. Novel polymeric nanoparticles for intracellular delivery of peptide Cargos: antitumor efficacy of the BCL-2 conversion peptide NuBCP-9. Cancer Res. 2014;74: 3271–81. doi:10.1158/0008-5472.CAN-13-2015

15. Fleishman SJ, Whitehead TA, Ekiert DC, Dreyfus C, Corn JE, Strauch E-M, et al. Computational design of proteins targeting the conserved stem region of influenza hemagglutinin. Science. 2011;332: 816–21. doi:10.1126/science.1202617

16. Berger S, Procko E, Margineantu D, Lee EF, Shen BW, Zelter A, et al. Computationally designed high specificity inhibitors delineate the roles of BCL2 family proteins in cancer. Elife. 2016;5. doi:10.7554/eLife.20352

17. Roberts KE, Cushing PR, Boisguerin P, Madden DR, Donald BR. Computational design of a PDZ domain peptide inhibitor that rescues CFTR activity. PLOS Comput Biol. 2012;8: e1002477. doi:10.1371/journal.pcbi.1002477

18. Chevalier A, Silva D-A, Rocklin GJ, Hicks DR, Vergara R, Murapa P, et al. Massively parallel de novo protein design for targeted therapeutics. Nature. Nature Publishing Group; 2017;550: 74–79. doi:10.1038/nature23912

19. Arkadash V, Yosef G, Shirian J, Cohen I, Horev Y, Grossman M, et al. Development of High Affinity and High Specificity Inhibitors of Matrix Metalloproteinase 14 through Computational Design and Directed Evolution. J Biol Chem. 2017;292:

20. Feng X, Barth P. A topological and conformational stability alphabet for multipass membrane proteins. Nat Chem Biol. 2016;12: 167–173. doi:10.1038/nchembio.2001

21. Debartolo J, Dutta S, Reich L, Keating AE. Predictive Bcl-2 family binding models rooted in experiment or structure. J Mol Biol. Elsevier Ltd; 2012;422: 124–144. doi:10.1016/j.jmb.2012.05.022

22. DeBartolo J, Taipale M, Keating AE. Genome Wide Prediction and Validation of Peptides That Bind Human Prosurvival Bcl-2 Proteins. PLOS Comput Biol. 2014;10: e1003693. doi:10.1371/journal.pcbi.1003693

23. Fernandez-Fuentes N, Oliva B, Fiser A. A supersecondary structure library and search algorithm for modeling loops in protein structures. Nucleic Acids Res. 2006;34: 2085–97. doi:10.1093/nar/gkl156

24. Mackenzie CO, Grigoryan G. Protein structural motifs in prediction and design. Curr Opin Struct Biol. 2017;44: 161–167. doi:10.1016/j.sbi.2017.03.012

25. Vanhee P, Verschueren E, Baeten L, Stricher F, Serrano L, Rousseau F, et al. BriX: A database of protein building blocks for structural analysis, modeling and design. Nucleic Acids Res. 2011;39: 435–442. doi:10.1093/nar/gkq972

26. Jacobs TM, Williams B, Williams T, Xu X, Eletsky A, Federizon JF, et al. Design of structurally distinct proteins using strategies inspired by evolution. Science. 2016;352: 687–90. doi:10.1126/science.aad8036

27. Mackenzie CO, Zhou J, Grigoryan G. Tertiary alphabet for the observable protein structural universe. Proc Natl Acad Sci U S A. 2016; 201607178. doi:10.1073/pnas.1607178113

28. Zheng F, Grigoryan G. Sequence statistics of tertiary structural motifs reflect protein stability. PLoS One. 2017;12: 1–25. doi:10.1371/journal.pone.0178272

29. Zhou Z, Grigoryan G. A general-purpose protein design framework based on mining sequence structure relationships in known protein structures. 2018; BioRxiv doi: https://doi.org/10.1101/431635.

30. Zheng F, Zhang J, Grigoryan G. Tertiary structural propensities reveal fundamental sequence/structure relationships. Structure. Elsevier Ltd; 2015;23: 961–971. doi:10.1016/j.str.2015.03.015

31. Opferman JT. Attacking cancer’s Achilles heel:antagonism of anti-apoptotic BCL-2 family members. FEBS J. 2016;283: 2661–75. doi:10.1111/febs.13472

32. Hiraki M, Maeda T, Mehrotra N, Jin C, Alam M, Bouillez A, et al. Targeting MUC1-C suppresses BCL2A1 in triple-negative breast cancer. Signal Transduct Target Ther. 2018;3: 13. doi:10.1038/s41392-018-0013-x

33. Souers AJ, Leverson JD, Boghaert ER, Ackler SL, Catron ND, Chen J, et al. ABT-199, a potent and selective BCL-2 inhibitor, achieves antitumor activity while sparing platelets. Nat Med. 2013;19: 202–8. doi:10.1038/nm.3048

34. Cang S, Iragavarapu C, Savooji J, Song Y, Liu D. ABT-199 (venetoclax) and BCL-2 inhibitors in clinical development. J Hematol Oncol. 2015;8: 129. doi:10.1186/s13045-015-0224-3

35. Foight GW, Ryan JA, Gullá S V., Letai A, Keating AE. Designed BH3 peptides with high affinity and specificity for targeting Mcl-1 in cells. ACS Chem Biol. 2014;9: 1962–1968. doi:10.1021/cb500340w

36. Jenson JM, Ryan JA, Grant RA, Letai A, Keating AE. Epistatic mutations in PUMA BH3 drive an alternate binding mode to potently and selectively inhibit anti-apoptotic Bfl-1. Elife. 2017;6: 1–23. doi:10.7554/eLife.25541

37. Berger S, Procko E, Margineantu D, Lee EF, Shen BW, Zelter A, et al. Computationally designed high specificity inhibitors delineate the roles of BCL2 family proteins in cancer. Elife. 2016;5: 1–31. doi:10.7554/eLife.20352

38. Kotschy A, Szlavik Z, Murray J, Davidson J, Maragno AL, Le Toumelin-Braizat G, et al. The MCL1 inhibitor S63845 is tolerable and effective in diverse cancer models. Nature. 2016;538: 477–482. doi:10.1038/nature19830

39. Jenson JM, Xue V, Stretz L, Reich L, Keating AE. Peptide design by optimization on a data parameterized protein interaction landscape. Proc Natl Acad Sci. in press 2018; doi:10.1073/pnas.1812939115

40. Reich L, Dutta S, Keating AE. SORTCERY - A High-Throughput Method to Affinity Rank Peptide Ligands. J Mol Biol. 2015;427: 2135–2150. doi:10.1016/j.jmb.2014.09.025

41. Lewis SM, Kuhlman BA. Anchored design of protein-protein interfaces. PLoS One. 2011;6: e20872. doi:10.1371/journal.pone.0020872

42. Alford RF, Leaver-Fay A, Jeliazkov JR, O’Meara MJ, DiMaio FP, Park H, et al. The Rosetta All-Atom Energy Function for Macromolecular Modeling and Design. J Chem Theory Comput. 2017;13: 3031–3048. doi:10.1021/acs.jctc.7b00125

43. Schymkowitz J, Borg J, Stricher F, Nys R, Rousseau F, Serrano L. The FoldX web server: An online force field. Nucleic Acids Res. 2005;33: W382–W388. doi:10.1093/nar/gki387

44. Fire E, Gullá S V, Grant RA, Keating AE. Mcl-1-Bim complexes accommodate surprising point mutations via minor structural changes. Protein Sci. 2010;19: 507–19. doi:10.1002/pro.329

45. Miles JA, Yeo DJ, Rowell P, Rodriguez-Marin S, Pask CM, Warriner SL, et al. Hydrocarbon constrained peptides - understanding preorganisation and binding affinity. Chem Sci. 2016;7: 3694–3702. doi:10.1039/c5sc04048e

46. Gai SA, Wittrup KD. Yeast surface display for protein engineering and characterization. Curr Opin Struct Biol. 2007;17: 467–73. doi:10.1016/j.sbi.2007.08.012

47. Dutta S, Gullá S, Chen TS, Fire E, Grant RA, Keating AE. Determinants of BH3 Binding Specificity for Mcl-1 versus Bcl-xL. J Mol Biol. 2010;398: 747–762. doi:10.1016/j.jmb.2010.03.058

48. Dutta S, Chen TS, Keating AE. Peptide ligands for pro-survival protein Bfl-1 from computationally guided library screening. ACS Chem Biol. 2013;8: 778–88. doi:10.1021/cb300679a

49. Jenson JM, Ryan JA, Grant RA, Letai A, Keating AE. Epistatic mutations in PUMA BH3 drive an alternate binding mode to potently and selectively inhibit anti-apoptotic Bfl-1. Elife. 2017;6. doi:10.7554/eLife.25541

50. Lee EF, Fedorova A, Zobel K, Boyle MJ, Yang H, Perugini MA, et al. Novel Bcl-2 homology-3 domain-like sequences identified from screening randomized peptide libraries for inhibitors of the prosurvival Bcl-2 proteins. J Biol Chem. 2009;284: 31315–31326. doi:10.1074/jbc.M109.048009

51. Zheng F, Jewell H, Fitzpatrick J, Zhang J, Mierke DF, Grigoryan G. Computational design of selective peptides to discriminate between similar PDZ domains in an oncogenic pathway. J Mol Biol. 2015;427: 491–510. doi:10.1016/j.jmb.2014.10.014

52. Davey JA, Chica RA. Improving the accuracy of protein stability predictions with multistate design using a variety of backbone ensembles. Proteins Struct Funct Bioinforma. 2014;82: 771–784. doi:10.1002/prot.24457

53. Procko E, Berguig GY, Shen BW, Song Y, Frayo S, Convertine AJ, et al. A computationally designed inhibitor of an Epstein-Barr viral Bcl-2 protein induces apoptosis in infected cells. Cell. Elsevier Inc.; 2014;157: 1644–1656. doi:10.1016/j.cell.2014.04.034

54. Kingsford CL, Chazelle B, Singh M. Solving and analyzing side-chain positioning problems using linear and integer programming. Bioinformatics. 2005;21: 1028–1039. doi:10.1093/bioinformatics/bti144

55. Grigoryan G, Reinke AW, Keating AE. Design of protein-interaction specificity gives selective bZIP-binding peptides. Nature. Nature Publishing Group; 2009;458: 859–864. doi:10.1038/nature07885

56. Chaudhury S, Lyskov S, Gray JJ. PyRosetta: a script-based interface for implementing molecular modeling algorithms using Rosetta. Bioinformatics. 2010;26: 689–91. doi:10.1093/bioinformatics/btq007

57. Crooks GE, Hon G, Chandonia J-M, Brenner SE. WebLogo: a sequence logo generator. Genome Res. 2004;14: 1188–90. doi:10.1101/gr.849004

58. Muñoz V, Serrano L. Development of the multiple sequence approximation within the AGADIR model of alpha-helix formation: comparison with Zimm-Bragg and Lifson-Roig formalisms. Biopolymers. 1997;41: 495–509. doi:10.1002/(SICI)1097-0282(19970415)41:5<495::AID-BIP2>3.0.CO;2-H

59. Gautier R, Douguet D, Antonny B, Drin G. HELIQUEST: a web server to screen sequences with specific alpha-helical properties. Bioinformatics. 2008;24: 2101–2. doi:10.1093/bioinformatics/btn392

60. Larkin MA, Blackshields G, Brown NP, Chenna R, McGettigan PA, McWilliam H, et al. Clustal W and Clustal X version 2.0. Bioinformatics. 2007;23: 2947–8. doi:10.1093/bioinformatics/btm404

61. Henikoff S, Henikoff JG. Amino acid substitution matrices from protein blocks. Proc Natl Acad Sci U S A. 1992;89: 10915–10919. doi:10.1073/pnas.89.22.10915

62. UniProt Consortium T. UniProt: the universal protein knowledgebase. Nucleic Acids Res. 2018;46: 2699. doi:10.1093/nar/gky092

63. Berman HM. The Protein Data Bank. Nucleic Acids Res. 2000;28: 235–242. doi:10.1093/nar/28.1.235

64. Needleman SB, Wunsch CD. A general method applicable to the search for similarities in the amino acid sequence of two proteins. J Mol Biol. 1970;48: 443–53.

65. McConkey BJ, Sobolev V, Edelman M. Discrimination of native protein structures using atom-atom contact scoring. Proc Natl Acad Sci U S A. 2003;100: 3215–20. doi:10.1073/pnas.0535768100

66. Wang S, Peng J, Xu J. Alignment of distantly related protein structures: Algorithm, bound and implications to homology modeling. Bioinformatics. 2011;27: 2537–2545. doi:10.1093/bioinformatics/btr432

67. Otwinowski Z, Minor W. [20] Processing of X-ray diffraction data collected in oscillation mode. Methods Enzymol. 276: 307–326. doi:10.1016/S0076-6879(97)76066-X

68. McCoy AJ, Grosse-Kunstleve RW, Adams PD, Winn MD, Storoni LC, Read RJ. Phaser crystallographic software. J Appl Crystallogr. 2007;40: 658–674. doi:10.1107/S0021889807021206

69. Emsley P, Lohkamp B, Scott WG, Cowtan K. Features and development of Coot. Acta Crystallogr D Biol Crystallogr. 2010;66: 486–501. doi:10.1107/S0907444910007493

70. Czabotar PE, Lee EF, Thompson G V, Wardak AZ, Fairlie WD, Colman PM. Mutation to Bax beyond the BH3 domain disrupts interactions with pro-survival proteins and promotes apoptosis. J Biol Chem. 2011;286: 7123–31. doi:10.1074/jbc.M110.161281

